# The sensorimotor basis of subjective experience in social synchronization behavior

**DOI:** 10.1101/2025.02.09.637302

**Authors:** Jan P. Bremer, Marius S. Knorr, Florian Göschl, Alexander Maye, Andreas K. Engel, Till R. Schneider

## Abstract

The sensorimotor contingency (SMC) theory addresses action and perception as constitutive factors to one another – what is being perceived depends on what we do and vice versa. Accordingly, perception is seen as an active process of probing the environment and receiving feedback from it. These action-effect patterns may also be a predominant factor in social interactions. They can manifest in the phenomenon of synchronization, for example, when applause, gait, or posture in conversations synchronize unintentionally and form the basis of the concept of socially deployed sensorimotor contingencies (socSMCs). In this study, we introduce and compare measures used to study complex systems in order to quantify the information-theoretic basis of socSMCs. Two human participants had to synchronize arm movements in a full body virtual reality (VR) environment with each other. We aimed to evaluate the information-theoretic measures transfer entropy and mutual information in this complex motion synchronization task. Furthermore, in our experiment participants shared a mutual sensation of synchronicity and creativity for their interaction solely based on their movements. These subjective ratings of synchronicity and creativity can be predicted using transfer entropy and mutual information, showing that informational coupling between agents is relevant to subjective experience of interaction as described by the concept of socSMCs.

## 2. Introduction

Classical cognitive science theories explain cognition and perception as the results of internal representations of the external world. These internal representations then lead to actions. However, recently there has been a shift in cognitive sciences towards more enactive action-oriented approaches to cognition and perception such as the sensorimotor contingency (SMC) theory (O ‘Regan and Noë, 2001; Noë, 2010, O ‘Regan, 2011; O ‘Regan and Noë, 2001). In contrast to classical frameworks, the theory describes perception and cognition as the result of probing the environment with our actions and perceiving its effects. These effects then form the basis for new actions – action and perception are thus influencing each other bidirectionally (Engel et al., 2013; Kaspar et al., 2014; König et al., 2013). According to the SMC theory, perception and cognition are understood as active processes based on these action-effect patterns (O ‘Regan and Noë, 2001).

The SMC theory can be extended to social interactions between biological and/or artificial agents. Accordingly, complex social interactions are viewed as being grounded in continuous sensorimotor patterns (Lübbert et al., 2021). These dependencies are described by the concept of socially relevant sensorimotor contingencies (socSMCs), which expands the classic SMC theory by applying it to the domain of social interactions (De Jaegher et al., 2010; Maye and Engel, 2016; Lübbert et al., 2021). Sensorimotor interactions are thought to be the fundamentals that provide knowledge about the intentions, beliefs, and personalities of other agents. As in the SMC theory, the building of social sensorimotor contingencies relies on the interaction between perception and action. These action-effect contingencies can explain behavior in everyday life situations. For example, in conversations we intuitively mirror our counterparts ‘ movements, facial expressions, or postures. Individuals in groups synchronize their movements in rituals, while dancing, or during applause and often they harmonize unintentionally during joint action.

The socSMCs concept suggests that sensorimotor contingencies are essential for the effective coupling of agents during social interactions (Lübbert et al., 2021). It describes that informational and sensorimotor coupling is the basis of successful social interactions, which can be described at three different levels of coupling between social agents: unidirectional information sharing (check SMCs), bidirectional coupling (sync SMCs), and multidirectional coupling (unite SMCs). Check SMCs involve unidirectional coupling with one agent sharing information with another one. Check SMCs may lead to social entrainment which in turn can result in sync SMCs that are based on bidirectional coupling between both agents sharing information and grounding their decisions and sensorimotor interaction on this shared information. This informational coupling may enable agents to predict the decisions of their counterpart. Unite SMCs implicate coupling between multiple agents, e.g. emergence of joint concepts or group habits, which are not in the focus of the present study.

In this study, we aim to investigate sensorimotor and informational coupling by defining two interacting agents as a single system using measures of information transfer to quantify the degree of coupling between these agents (Fusaroli and Tylén, 2016). Here, we use transfer entropy as a measure which determines information flow between two processes to quantify the directed interaction between two social agents (Orange and Abaid, 2015) and mutual information, a measure which serves to capture shared information between processes, to quantify shared information between interacting agents.

Recently, these information-theoretic measures received a lot of interest in neuroscience (Lizier et al., 2010; Vincente et al. 2010). Transfer entropy is traditionally used to estimate the directional information transfer between two coupled processes. Mutual information, on the other hand, is a measure to quantify the uncertainty reduction of one process given information of the coupled process and has been used successfully to measure inter-brain coupling in social interactions (Naeem et al. 2012).

In this study, we utilize a motion synchronization task to investigate the information-theoretic basis of socSMCs. Recently, a series of experiments investigated social interactions using the mirror game (Noy et al., 2011), a motion synchronization task based on improvisation in theater and music, in which the degree of synchronization between agents can be quantified. The mirror game serves as a paradigm for investigating synchronization of movements and joint improvisation, in the original study by Noy and colleagues (2011) with simple one-dimensional hand movements. Two participants had the task to synchronize their hand movements, while holding handles and moving them on a given trajectory. Participants were instructed to either lead or follow the movements of their counterpart or to jointly improvise movements without designated leader or follower roles. Noy and colleagues (2011) showed that during improvisation participants produced more complex and synchronous movements. A more recent study investigated the mirror game in a naturalistic and complex two-dimensional scenario and reported the subjective sensation of synchronization to be related to performance parameters derived from participants ‘ movements (Llobera et al., 2016). These findings emphasize a substantial interdependence of action and perception in social interactions. In the current study, we investigate informational and sensorimotor coupling of movement synchronization behavior and hypothesize increased bidirectional coupling during improvisation compared to instructed behavior.

In the classical mirror game, participants are instructed to take one of three roles when interacting: leading, following, or jointly improvising. These roles are based on social hierarchies. Interactions can have a hierarchical structure that results in leader and follower roles, but real-life situations often involve joint action and improvisation without a designated leader or follower: for example, when people have to synchronize their actions to jointly lift an object, movement information has to be shared bidirectionally. Hierarchical structures can also emerge in improvisation scenarios and can be dependent on countless observable factors, such as appearance, gender, voice and many more. Visualizing the own and the counterpart ‘s bodies only as neutral avatars in a virtual environment and thereby reducing the human body to its movements is essential for investigating motion synchronization without confounding factors. We developed a virtual reality (VR) setup which uses 3-dimensional, naturalistic movements, to increase ecological validity and implications for real life compared to one-dimensional mirror games. This study is, to the best of our knowledge, the first to perform a three-dimensional movement synchronization task with movement quantification in VR.

## 3. Materials and Methods

### 3.1. Participants

A total of 50 participants without any record of psychiatric or neurological illnesses were recruited and randomly assigned to 25 pairs of two players, randomly mixing male and female participants. Participants of each pair arrived independently of each other and did not meet before or during the experiment. One of the 25 pairs had to be excluded from the analysis because one participant interrupted the experiment due to illness. The final sample consisted of 48 participants (aged 19 – 33 years; mean = 22.79 years, standard deviation (SD) = 2.73; 28 female and 20 male) who were eligible for the analyses. Participants gave written consent and were financially compensated for their participation. The study was approved by the local ethics committee of the Medical Association in Hamburg, Germany, and adhered to the Declaration of Helsinki.

### 3.2. Hardware

The participants ‘ motion was tracked by two Xsens Motion Tracking Systems (MTw Awinda, Xsens, The Netherlands) with a 60 Hz sampling rate using eleven sensors on the upper body, which were placed unilaterally on head, pelvis, sternum and bilaterally on shoulders, hands, upper and lower arms. Two identical virtual environments were created in Unity3D (Unity Technologies, USA) and connected via TCP sockets. Each environment was screened on an HTC Vive Pro VR-System (HTC/Valve, Taiwan) driven by a GeForce RTX 2080 graphics card (NVIDIA, USA).

### 3.3 Software

Motion trajectories were broadcasted in real-time and rendered via Unity3D to each VR-System using the HTC Vive system. Mutual information and transfer entropy were calculated with the Java Information Dynamics Toolkit (JIDT) in Python (Lizier, 2014). The repeated measures correlations calculation of felt synchronicity, creativity, sympathy and focus between both participants were calculated using pingouin (v. 0.3.3) a statistical package for Python. For the generalized linear-mixed model analysis SPSS 27 (IBM, USA) was used.

### 3.4. Experimental procedure

Participants of each pair were tested in different rooms and did not meet until the end of the experiment. They were not informed in any way about their partner beforehand. Participants received detailed instructions about the procedure both orally and in written form. Before the movement synchronization task started, participants completed the following personality questionnaires: autism quotient (AQ; Baron-Cohen et al., 2001), SPF (German version of the Interpersonal Reactivity Index, IRI; Davis, 1983; Paulus, 2009), and NEO-FFI (Costa and McCrae, 1992). The AQ is a self-report inventory aiming to test for symptoms of autism spectrum conditions. The SPF is a self-report questionnaire investigating empathic personality traits. And the NEO-FFI is a short version of the NEO PI-R (personality inventory) examining the Big Five personality traits (openness to experience, conscientiousness, extraversion, agreeableness, and neuroticism). Additionally, participants were asked to complete a customised questionnaire about their current physical well-being and regarding their prior experience with VR scenarios, musical instruments, and dancing.

Participants were equipped with the Xsens motion tracking system. After the calibration of the motion tracking system, participants put on the VR goggles. The functionality and calibration of the motion tracking and the VR system were checked by the experimenter. At this moment, participants had the possibility to gain a first impression and get used to the VR environment. Participants were able to see the movements of their avatar in first person (Figure 1a). Movements of both participants were instantaneously projected onto two avatars without noticeable delay. At this point the two avatars were visually separated using a virtual wall and, thus, participants could only see parts of their own avatar. This procedure led to the impression that the avatar was fully controlled by the participant ‘s own movements. As soon as both participants - situated in two different rooms - were ready, the experiment started.

**Figure 1.**
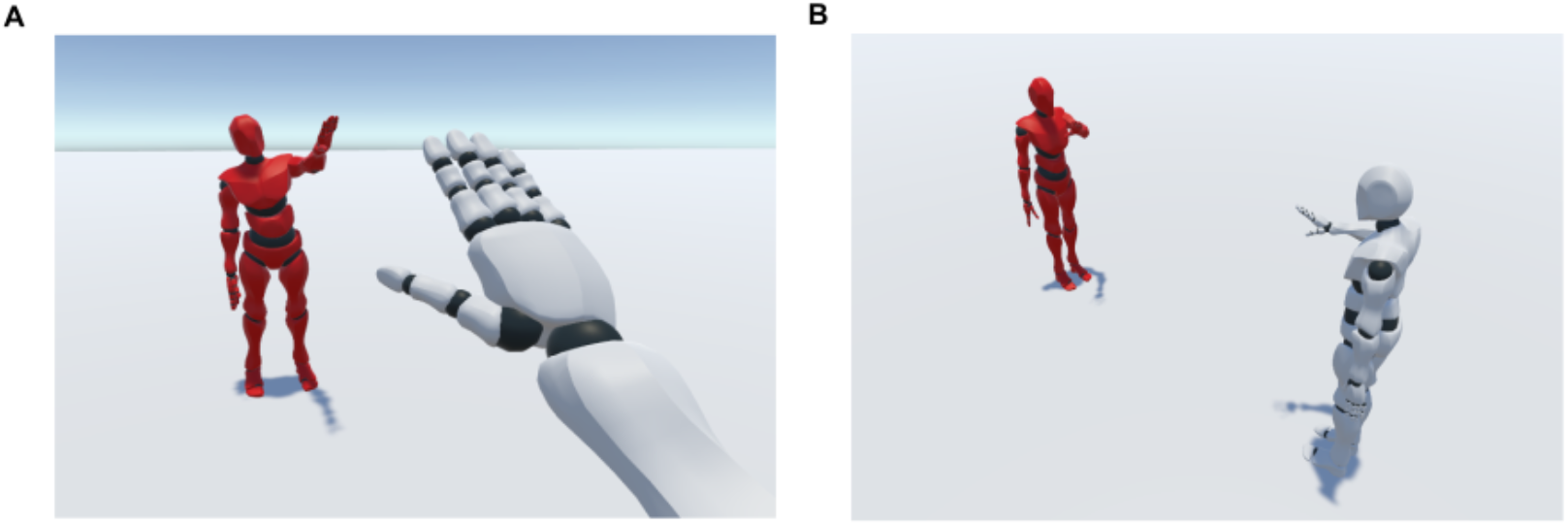
Virtual reality setup. **A**. Perspective of the scene view in the HTC Vive system (white: the own avatar, red: the other participant ‘s avatar). This is the scene that participants were seeing during the experiment. For both participants the partner was displayed in red. **B**. Bird ‘s eye view of the game (not visible for participants).

In the virtual environment, two avatars were visible standing face-to-face to each other. The perspective of each participant was egocentric, as if the avatar body was one ‘s own body. Both partners were instructed to synchronize their arm movements. One partner was instructed to use the right arm and the other partner the left arm, so that mirror movements were possible in the virtual setup (Figure 1a, b). There were no restrictions concerning the nature of arm movement. Partners were instructed to focus the synchronization task only on the arm movement and to avoid eccentric movements of the remaining limbs.

Before each trial, participants were instructed to take one of three different roles. They could either lead (L), follow (F) or jointly improvise (JI) arm movements. The leader was instructed to produce arm movements which the partner should follow. In the joint improvisation task, partners were instructed to synchronize their movements without a designated leader or follower. The experiment consisted of 15 trials lasting 2 min each. The total interaction time was 37 min. The order of conditions was L, F, JI – JI, L, F – L, F, JI – L, F, JI – JI, F, L. Between trials there was a pause of 50 seconds in which partners were asked to rate their experience during the last trial regarding focus, synchronicity, creativity and sympathy for their counterpart on a continuous scale (range 1-10). A short resting period followed in which they received the instruction for the next trial.

The experiment was followed up with a questionnaire regarding the experience of the interaction. Participants rated their perceived role in the joint improvisation condition on an ordinal scale (range 1-10, 1: Leader; 10: Follower) and whether the interaction consisted of co-leadership or changing leader/follower roles (range 1-10, 1: co-leadership, 10: changing leader/follower roles).

### 3.5. Data generation and preprocessing

Motion data from 48 participants interacting in leader, follower and joint improvisation conditions was measured (5 trials per condition, each lasting 2 min) resulting in 360 trials in total (24 pairs acting in 15 trials each, 240 trials leader/follower, 120 trials joint improvisation). The data consisted of eleven quaternions describing the relative motion provided by the eleven sensors on the upper body. Quaternions are 4-dimensional representations of rotations in 3-dimensional space (w, x, y, z) and represent the rotation of an angle (α) around the axis of a vector from the origin (0,0,0) to (x,y,z). From this the quaternion is derived as:

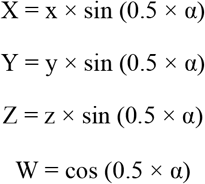

The quaternions are mapped to a uniform avatar independent of the participants body dimensions in Unity. Therefore, further analysis was independent of the participants body dimensions. The origin of the coordinate system was the pelvis center. Further analyses were limited to the three-dimensional trajectories of the active hand. The data was recorded at 60 Hz. The data was filtered to remove any jitter in the movement using a Savitzky-Golay filter (polynomial of third degree, window length of 17, which represents 0.28s at a sampling frequency at 60Hz). Only one of 360 trials was excluded due to technical failure (n = 1).

### 3.6. Transfer entropy

Transfer entropy (Schreiber, 2000, Barnett et al., 2009) is a non-parametric measure, which represents the directional transfer of information between two processes by calculating the reduction of uncertainty with the current information of one system in predicting the future state of the other system. Transfer entropy is measured in bits, the basic unit of information.

Here we use transfer entropy to calculate the flow of information from one participant ‘s hand movements to the other participant ‘s hand movements. In a movement-based leader-follower scenario, information from the leader is transferred to the follower, influencing the follower ‘s future movements. If information of the leader ‘s movement can be used to predict the future movements of the follower, transfer entropy from leader to follower is larger than transfer entropy from follower to leader. Accordingly, this method can be used to classify leaders and followers in corresponding scenarios as well as in open-end improvisation. In contrast, an improvisation task should require a more balanced exchange of information between both parties. Here, transfer entropy may determine leader and follower tendencies.

We computed transfer entropy on continuous data using the Kraskov-Stogbauer-Grassberger transfer entropy estimator and searched for the maximum transfer entropy in a search space of 1500 ms on each individual axis of movement (x,y,z) per trial (Figure 2). We tested the transfer entropy between conditions using paired t-tests (Table 1). Furthermore, given the directional nature of transfer entropy, we calculated net transfer entropy (netTE) (Figure 3):

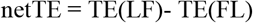

**Figure 2.**
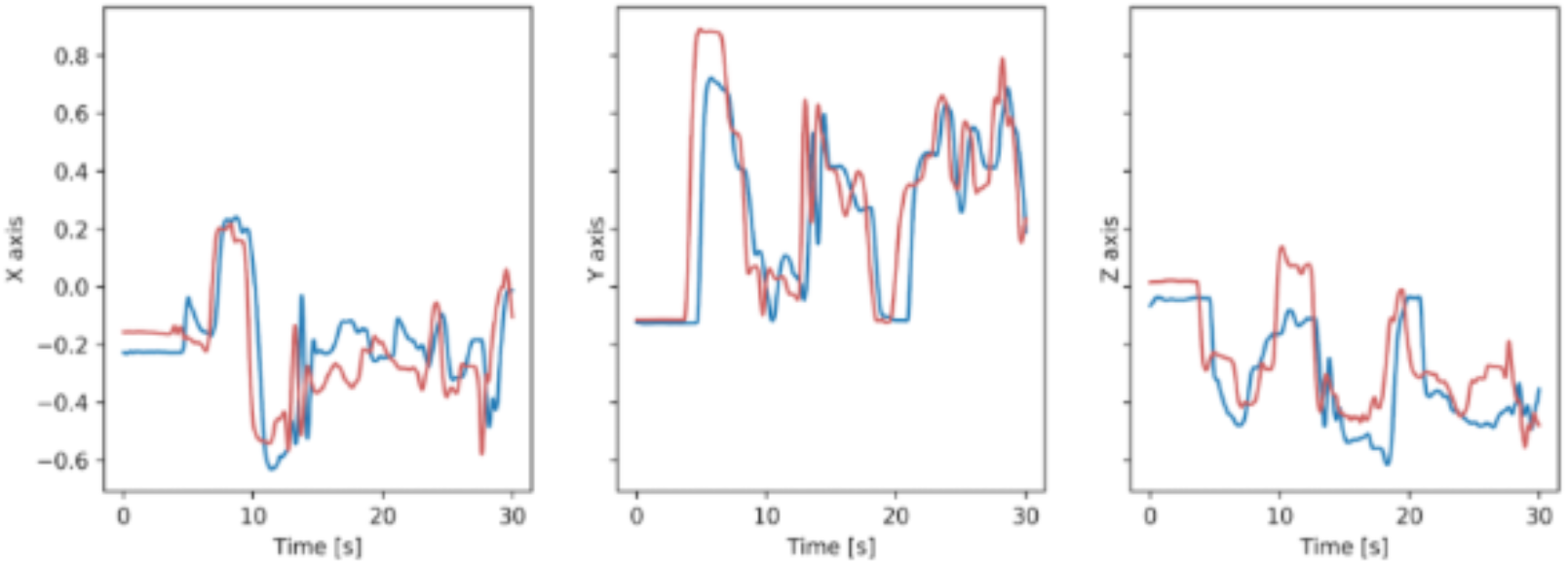
Example of leader (red line) and follower (blue line) trajectories showing synchronized movements in three axes (x, y, z) in a leader/follower trial.

**Table 1.** Statistical evaluation of transfer entropy between follower and leader (F), leader and follower (L), joint improvisation (JI). Only the comparison between JI and F conditions shows significant results for the y-axis (df =23; p < 0.05, Bonferroni corrected)

| Axis | F-L | JI-L | JI-F |
| --- | --- | --- | --- |
| x-axis | $t = 0.334, p = 0.739$ | $t = 0.4291, p = 0.669$ | $t = 0.8307, p = 0.410$ |
| y-axis | $t = 1.67, p = 0.101$ | $t = 1.285, p = 0.204$ | $t = 3.0941, p = 0.003$ |
| z-axis | $t = 0.4741, p = 0.637$ | $t = 0.7054, p = 0.483$ | $t = 0.2085, p = 0.835$ |

**Figure 3.**
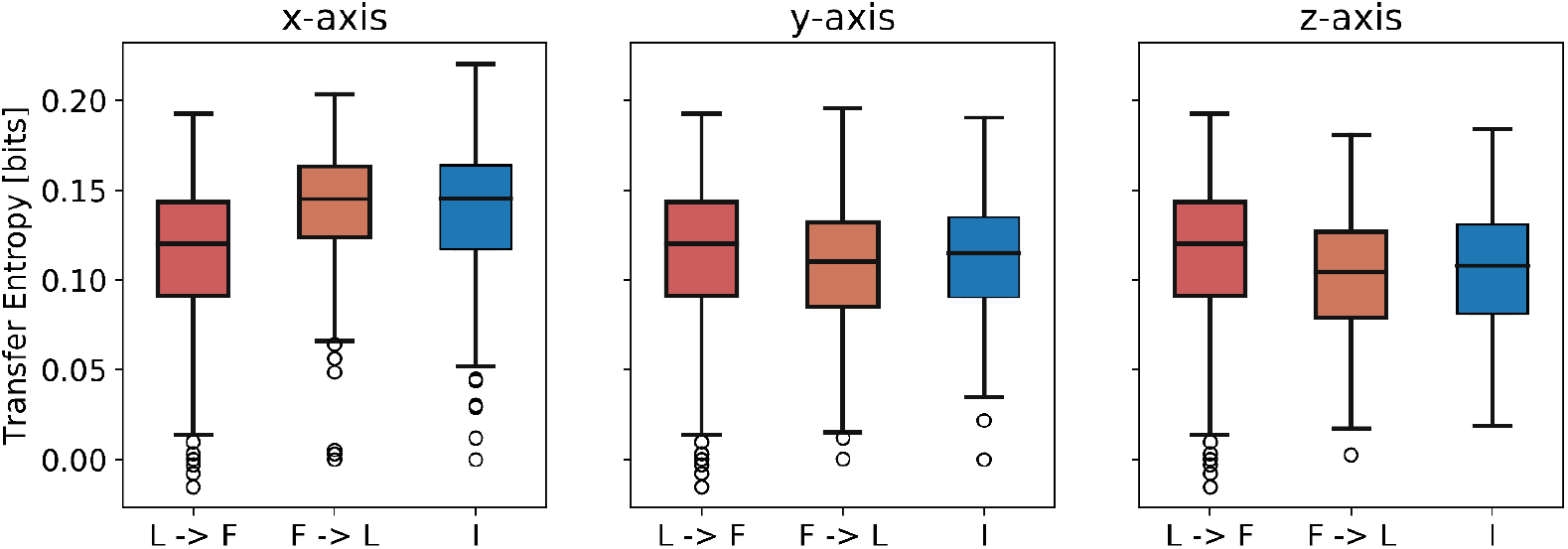
Transfer entropy [bits] from follower to leader, from leader to follower, and in JI conditions for the three spatial axes. Medians with interquartile ranges are shown (outliers being represented as circles).

Based on the assumption that information flow from leader to follower should be greater than vice versa, netTE should be positive in leader-follower scenarios. We tested this netTE from leader to follower using t-tests against a normal distribution with a mean of zero (Nakayama et al., 2017).

### 3.7. Mutual information

Mutual information is a symmetric statistic that quantifies the amount of information that two processes have in common. Mutual information represents the reduction of uncertainty about one process by learning from the other process and vice versa. In a motion synchronization task mutual information represents the information both participants base their consecutive movements on. We use this as a measure of bidirectional coupling and synchronicity. In improvisation scenarios, participants have to rely on their shared information of the process to conclude further action. In contrast, leader-follower scenarios do not rely on the bi-directional exchange of information as heavily as improvisation tasks. To calculate mutual information, we used the Java Information Dynamic Toolkit ‘s (JIDT) implementation of mutual information on continuous data, also utilizing the Kraskov-Stogbauer-Grassberger estimator.

### 3.8. Task performance parameters

Similar to the original mirror game (Noy, et al., 2011) we calculated the motion velocity and velocity error. Motion velocity was assessed by calculating the mean of the three-dimensional Euclidean distance per time point of both participants in 3D space as:

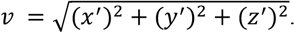

and mean velocity as:

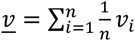

The velocity error was assessed as the absolute difference in velocities per time point.

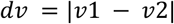

and mean velocity error as:

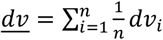

### 3.9. Statistical methods

Using the GENLINMIXED package in SPSS, we deployed a generalized linear mixed effect model to determine whether net transfer entropy, transfer entropy, mutual information and velocity error are predictive of the ratings of synchronicity and creativity after each trial. A hierarchical linear model was built with trials on the lowest level, participants on the second level and pairs on the third level. To reduce the number of features for this model, we applied principal component analysis (PCA) on mutual information and transfer entropy to reduce the three axes of these measurements to the first principal component. In the linear mixed-model either synchronicity or creativity were used as outcome variable. Subject ID and Pair ID were considered as random effects, Trial ID as the repeated measure variable. A backward elimination procedure was used for feature selection. In the backward elimination process, the model was first fitted with all predictors (net transfer entropy, transfer entropy, mutual information and velocity error). Step-by-step the predictor with the highest p-value was removed from the model and the model was fitted again until only significant predictors and interactions remained.

## 4. Results

Participants performed movements varying in amplitude and frequency in the three axes (Figure 2 a-c) and synchronized with each other. Unlike in the original mirror game (Noy et al., 2011), we did not observe the characteristic jitter of the follower ‘s motion velocity around the leader ‘s motion velocity, but instead a smoother trajectory lagging behind the leader ‘s movement.

### 4.1. Task performance

Motion velocity was significantly lower in JI than in L/F (t(23) = - 4.8870, p < 0.001). As a measure of synchronization, we calculated the velocity error (see Noy et al., 2011), which was not significantly different in JI compared to L/F trials (t(70) = - 0.86409, p = 0.396455). Mutual information in all axes was significantly higher in JI than in L/F trials (x-axis: t(23) = - 3.577, p = 0.002; y-axis: t(23) = - 3.285, p = 0.003; z-axis: t(23) = - 3.684, p = 0.001), indicating that participants based their consecutive movements more on shared information in the JI than in the L/F condition.

### 4.2. Leader- and followership

In order to classify leader- and followership in JI without a designated leader we calculated transfer entropy as a per-trial-measure of leader- and followership (Table 1, Figure 3). Transfer entropy can be calculated from leader to follower (L -> F), follower to leader (F -> L) and between the participants in joint improvisation (JI). Transfer entropy was only significantly different between conditions in the y-axis, showing smaller values for joint improvisation compared to follower condition (t(23) = 3.0941, p = 0.0033)

Net transfer entropy in L/F was positive in 77,1%, 79,2%, 72,9% (x axis, y axis, z axis) of participant pairs (Figure 4) and differed significantly from zero between leader and follower (x-axis: t(23) = - 6.597, p < 0.001, y-axis: t(23) = - 5.6825, p < 0.001, z-axis: t(23) = - 7.1751, p < 0.001; Figure 4)

**Figure 4.**
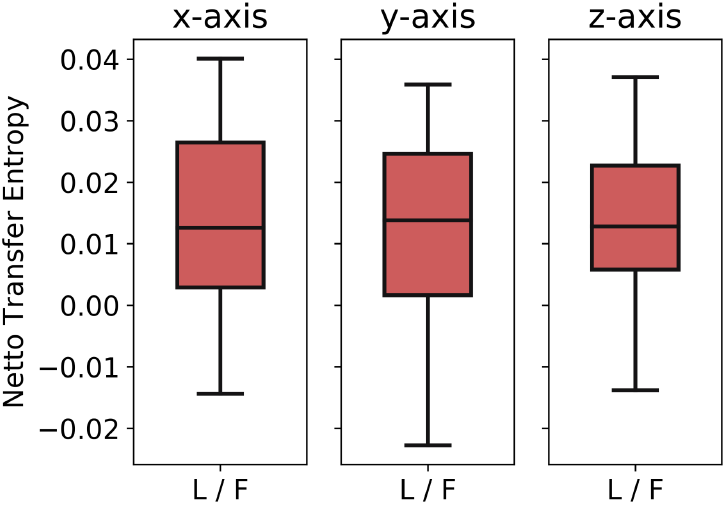
Net transfer entropy [bits] (leader - follower) for the three spatial axes.

### 4.3. Ratings

After each trial we assessed focus (Lead: Follow: JI: 8.29 +-1.48), experienced synchronicity (Lead: 7.32 +-1.73 Follow: 7.19 +-1.61, JI: 7.27 +-1.56), creativity (Lead: 7.20 +-1.78, Follow 6.55 +-1.73, JI: 6.85 +-1.88) and sympathy (Lead: 8.31 +-1.51, Follow: 8.38 +-1.41, JI: 8.55 +-1.33) We calculated the repeated measures correlation of synchronicity (r = 0.15, p = 0.002), sympathy (r = 0.076, p = 0.197), creativity (r = 0.116, p = 0.042), and focus (r = 0.11, p = 0.047) between pairs of participants.

We used a generalized linear mixed model to further investigate the relationship between the movement parameters and the ratings of perceived synchronicity and creativity of the participants. The analysis revealed that mutual information of the two participants is predictive of the perceived synchronicity (F(1,613) = 6.485, p = 0.011, β = 0.021; Table 2 and Supplementary Table 1). Furthermore, condition (F(2,608) = 3.322,p = 0.037) and the interaction of trial and condition are predictive of synchronicity (F(2,578) = 3.702, p = 0.025). Creativity ratings can be predicted by transfer entropy (F(1,431) = 10.65, p = 0.001, β = -0.022), (Table 3 and Supplementary Table 2), condition (F(2,527) = 15.305, p < 0.001).

**Table 2.** Results of the generalized linear mixed model analysis predicting synchronicity ratings revealing several fixed effects after backward elimination.

| Variable | F | p | $\beta$ |
| --- | --- | --- | --- |
| Mutual Information | 6.485 | 0.011 | 0.021 |
| Condition | 3.322 | 0.037 |  |
| Trial*Condition | 3.702 | 0.025 |  |

**Table 3.** Results of the generalized linear mixed model analysis predicting creativity ratings by several fixed effects after backward elimination

| Variable | F | p | $\beta$ |
| --- | --- | --- | --- |
| Transfer Entropy | 10.65 | 0.001 | -0.022 |
| Condition | 15.305 | < 0.001 |  |

In the post experimental inquiry, participants reported on their experience regarding follower- and leadership in the joint improvisation condition - whether the improvisation felt like switching between leader- and followership or a co-leadership in which participants were not aware of a designated leader. Participants rated on a scale from 1 (switching) to 10 (co-leadership) using a visual analogue scale (M = 4.12, SD = 2.87). When in a state of switching between leader- and followership, participants also reported whether they perceived themselves predominantly in the role of a follower or leader (rated on a scale from 1 (follower) to 10 (leader), M = 5.10, SD = 1.29). These results suggest that participants tend to perceive the improvisation condition as a switching between follower and leader instead of a co-leadership.

## 5. Discussion

This study has focused on the investigation of informational and sensorimotor coupling of human agents synchronizing their arm movements in a three-dimensional VR imitation task. In addition, the predictive value of these coupling measures on subjective experience of sensorimotor interactions was assessed. We instructed participants to interact in different scenarios and analyzed different information-theoretic measures, such as transfer entropy and mutual information. We showed that participants have a shared sensation, which can be partly predicted by coupling measures of their motion data. These results pave the way for future research investigating more open-ended, uninstructed interactions with dynamic coupling in social contexts. What is more, this study is the first to deploy human movement synchronization tasks in VR. For future studies, we propose this paradigm is suitable to investigate dynamic embodied interactions in more naturalistic scenarios.

### The mirror game

Noy and colleagues (2011) introduced the mirror game as a motion synchronization task using one-dimensional sliders with instructed leader/follower roles as well as improvisation. Over the years, this experimental design has been modified with a trend towards more naturalistic, real-life synchronization behavior. Recently, this development led to the deployment of a two-dimensional mirror game design (Llobera et al., 2016). Here, we developed the mirror game further into a three-dimensional version in a VR setup using motion tracking in combination with VR headsets.

In the original mirror game movements on one-dimensional sliders have shown a characteristic jitter of the follower around the leader ‘s trajectory. In our study this characteristic jitter was absent, potentially caused by the increased difficulty of the task compared to previous mirror game versions. Moreover, comparing improvisation and leader/follower conditions showed that motion velocity in improvisation was generally slower with smaller errors in velocity, in contrast to the original mirror game, in which participants ‘ interactions were faster with smaller velocity error (Noy et al., 2011). This may be due to the difference that Noy and colleagues studied experts in improvisation and used a less complex one-dimensional task in comparison to the free arm movements participants were able to perform in our experiment.

### Socializing sensorimotor contingencies

Building on the concept of socially deployed sensorimotor contingencies (socSMCs) Lübbert and colleagues (2021) distinguish three different stages of human interactions: sync SMCs, check SMCs and unite SMCs. Of these stages, only sync SMCs and check SMCs are relevant in this study. Firstly, check SMCs involve unidirectional coupling: agents determine how their counterpart reacts and can form predictions about their counterpart ‘s actions. Secondly, sync SMCs involve bidirectional coupling, which can lead to entrainment of behavior and shared action goals. Here we lay the foundation of quantifying behavioral coupling by comparing information-theoretic measures in instructed leader/follower role-taking mimicking check SMCs and improvisation mimicking sync SMCs.

To this end, we applied two information-theoretic measures for the analysis of human movement interactions. Firstly, we were able to quantify the information flow between leaders and followers using transfer entropy and net transfer entropy in instructed leader-follower interactions. While transfer entropy itself did not show significant differences between conditions, net transfer entropy was significantly different from zero, correctly classifying follower- and leadership. Thereby, we substantiate findings that leadership can be predicted by the direction of transfer entropy, as shown previously in studies involving leader-follower scenarios in animals (Sun et al., 2014; Orange and Abaid, 2015; Crosato et al., 2018) and speaker-to-listener scenarios in humans (Dumas et al., 2014). Furthermore, our findings on net transfer entropy substantiate findings of more restricted human interaction tasks to classify leader- and followership (Nakayama et al., 2017).

Secondly, we deployed mutual information, which is sought to quantify shared information, upon which the actions of the two participants are based. Mutual information values were significantly higher in the joint improvisation task condition than in conditions with well-defined leader and follower roles, indicating that the process of interacting is more strongly based on shared information in dynamic role-taking than in instructed role-taking.

### Subjective experience

Furthermore, we investigated whether participants ‘ shared sensation of synchronicity and creativity may be grounded in objective motion parameters and informational coupling in the movement synchronization task. Our results suggest that participants had a shared feeling of synchronicity and creativity that was potentially solely based on movement perception and generation as there was no verbal communication between participants possible. Therefore, we aimed to predict the sensation of synchronicity and creativity in our task from the information-theoretic measures using a generalized linear mixed model. Results show that mutual information is a significant predictor for the sensation of synchronicity. We interpret mutual information as the information both participants share before making a decision about their next move. High mutual information values were predictive of high synchronicity ratings. Velocity error on the other hand, a directly motion-dependent measure of synchronicity in the task, showed no predictive value.

For the sensation of creativity, the measure of transfer entropy showed significant predictive power. Our results revealed that higher transfer entropy is associated with one ‘s own leadership. This suggests that a stronger adoption of the leadership role by oneself is accompanied by a higher sensation of creativity.

### Limitations of the study

Three-dimensional motion synchronization is a complex task for the participants to perform and the analysis requires simplification. Thus, only hand movements were considered here but movements of other body parts were not considered. Even though we tried to keep the VR scenario as naturalistic as possible, participants may act differently compared to natural in-person synchronization. Most participants were young undergraduate students, future work should focus on a more diverse study sample.

## Conclusion

Our study demonstrates that social interactions can be studied in VR with embodied avatars. We used motion tracking in a two-person mirror game allowing 3-dimensional movements. The findings of our study support the notion that social interactions are grounded in modes of sensorimotor and informational coupling. These coupling modes, which are based on low-level sensorimotor interactions, can serve as a tool to predict subjective experience of social interactions. Transfer entropy has proven to successfully distinguish leader- and followership in this complex movement task. Transfer entropy and mutual information were predictors of the sensation of synchronicity and creativity, showing that shared information as well as the exchange of information are relevant for the experience of interaction. Furthermore, these findings have potential implications for the understanding of neuronal coupling in the human brain, which could be investigated using non-invasive methods such as electroencephalography (EEG). The neuronal foundations of social cognition and especially predictive coding could be investigated by combining the VR setup with the investigation of neuronal coupling between brain areas and between agents based on EEG signals.

## Acknowledgements

This work was supported by grants from the EU (project “socSMCs”, H2020-641321) and the DFG (SFB “Cross-Modal Learning”, TRR169-261402652-B1/Z2). We would like to thank Karin Reimann, Dept. of Neurophysiology and Pathophysiology, UKE Hamburg, for participant recruitment.

