## Supplementary material for "The sensorimotor basis of subjective experience in social synchronization behavior"

**Synchronicity**

| **Variable** | **F** | **p** |
|---|---|---|
| net Transfer Entropy | 0.046 | 0.83 |
| Transfer Entropy | 1.856 | 0.174 |
| Velocity Error | 2.49 | 0.115 |
| Mutual Information | 3.754 | 0.053 |
| Trial | 3.101 | 0.08 |
| Condition | 2.918 | 0.055 |
| Trial*Condition | 3.847 | 0.022 |

**Supplementary table 1.** Results of the generalized mixed linear model analysis without backwards elimination for synchrony.

**Creativity**

| **Variable** | **F** | **p** |
|---|---|---|
| net Transfer Entropy | 2.966 | 0.086 |
| Transfer Entropy | 9.718 | 0.002 |
| Velocity Error | 1.882 | 0.171 |
| Mutual Information | 0.155 | 0.694 |
| Trial | 0.248 | 0.619 |
| Condition | 3.489 | 0.031 |
| Trial*Condition | 1.698 | 0.184 |

**Supplementary table 2.** Results of the generalized mixed linear model analysis without backwards elimination for creativity.
